# Biofluorescence reveals hidden patterns in chitons with implications to visual ecology

**DOI:** 10.1101/2023.09.13.557364

**Authors:** Guido G. Grimaldi, Raiane dos Santos Guidi, Jaime A. Jardim, Marina Hirota, Daniel Marques Almeida Pessoa, Paulo Antunes Horta

## Abstract

Biofluorescence is apparently widespread in the tree of life. Fluorescence has the potential to contribute to the totality of light leaving an organism’s surface and will therefore circumscribe how an individual could be detected and discriminated by their visual predators. Here, we: (i) documented the first record of biofluorescence on polyplacophorans, (ii) spectrally characterized the biofluorescence on shells of living chitons, (iii) measured the colour patch pattern variation, (iv) separated the colour of their shells into their reflectance and fluorescence components, and (v) combined these data with field measurements to calculate perceptual distance in chromatic and achromatic contrasts based on the visual system of their major visually guided predators. We found a red biofluorescence that enhances the brightness of chiton shells, along with a correlation showing that as individuals grow larger, the fluorescing surface area becomes smaller. Our visual models suggest that fluorescence decreases the achromatic contrast of chitons against their naturally fluorescent substrates for most visual predators, making them less noticeable to specific predators. Our results support the potential visual functionality of biofluorescence and open new hypotheses regarding its ecological roles to further investigations.

## Introduction

The ability to perceive and respond to colours plays a significant role in the evolution of different species, shaping their visual systems, colour patterns, and behaviours in response to their environment and selective pressures (Osorio & Vorobyev, 1997; van der Kooi *et al*., 2021; Hagen *et al*., 2023). Many animals use colours for a range of competing functions, such as thermoregulation (Stuart-Fox *et al*., 2017), photoprotection (Lahti & Ardia, 2016), conspecific recognition (Siebeck *et al*., 2010), attracting mates (Endler & Houde, 1995), capturing preys (Heiling *et al*., 2003), and avoiding parasites (Caro *et al*., 2019) and predators (*e.g.,* camouflage, Price *et al*., 2019; *warning signals*, Lim *et al*., 2019; *mimicry*, Davis Rabosky *et al*., 2016; and *private communication*, Marshall & Stevens, 2014). However, colour perception is subjective and can be influenced by a variety of factors, including lighting conditions, surrounding colours (backgrounds), and differences in the spectral sensitivities of photoreceptor cells (Endler, 1978, 1993; Nilsson *et al*., 2022). By studying the visual systems of different animals, researchers can identify the spectral range of light that they can detect and use this information to create visual models that simulate the colour perception of different animals, allowing us to understand how they see the world around them, regardless of human experience (Kelber & Osorio, 2010; Renoult *et al*., 2017). Colours are typically described in terms of their hue, saturation, and brightness. As a result, organisms create chromatic and achromatic contrast against their background by using differences in colour (spectral reflectance patterns) and brightness (spectral intensity), both of which can significantly impact visual perception (Kelber *et al*., 2003).

Biological pigments and structural mechanisms are responsible for producing a wide variety of colours and patterns, by absorbing and reflecting specific wavelengths of light (Caro *et al*., 2017). Among them, biofluorescence compounds opens up a new range of visual possibilities, once it adds new photons to the totality of light that leave off a certain surface (exitance) (Marshall & Johnsen, 2017). This is possible because fluorescent pigments selectively absorb light in the short-wavelength range and re-emit it at a different one (Johnsen, 2011). So, the quantitative contribution of fluorescence photons to the exitance light depends on the environmental light and affect how animals can be seeing by their predators. Many living organisms exhibit biofluorescence, and it appears to be spread throughout the tree of life, as researchers have found it in vertebrates (Arnold, 2002; Lamb & Davis, 2020; Prötzel *et al*., 2021; Olson *et al*., 2021), invertebrates (Mazel, 2004; Lim *et al*., 2007; Eyal *et al*., 2015), and plants (Gandía-Herrero *et al*., 2005). Previous works bring potential visual functions to biofluorescence, such as mate choice (Arnold, 2002; Lim *et al*., 2007), pollinator (Thorp *et al*., 1975) and symbiotic algae attraction (Aihara *et al*., 2019), warning signals (Nemésio, 2005), fluorescent lures (Haddock, 2005), and camouflage (Sparks *et al*., 2014). Here, we add the first record of a biofluorescent chiton species and explore possible visual function of its fluorescence.

Chitons are polyplacophoran molluscs that possess a shell made up of eight overlapping plates or valves, which provide both flexibility and protection (Connors *et al*., 2012). They have a variety of sensory organs embedded throughout their shell that are connected by an intricate system of channels and pores filled by the chiton’s nervous system (Fernandez *et al*., 2007; Sumner-Rooney & Sigwart, 2018). This combination provides a unique and sophisticated sensory system that allows it to navigate, find food, avoid predators, and quickly and effectively respond to changes in its surrounding environment, making its shell an integrated sensory unit (Liu *et al*., 2022; Chappell & Speiser, 2023). Over evolutionary time, chitons evolved to occupy a variety of ecological niches, from shallow intertidal zones to deep sea environments (Saito *et al*., 2008; Gagnon *et al*., 2012; Sigwart, 2017; Mercegue *et al*., 2021). They play an important role in controlling algal growth which makes them an important part of many marine food webs (Littler *et al*., 1995; Bracken *et al*., 2018). Even though they are primarily herbivorous, several different organisms or even substratum particles may be found in their gut content (Sigwart & Schwabe, 2017). Intertidal chitons have a tidal activity rhythm, emerging during low tides to scrape food from the substratum and returning to refuge after each excursion period, but this seems to vary between species and location (Liversage & Benkendorff, 2017; Montecinos *et al*., 2020). Previous research has suggested that chitons may have their shell colours influenced by visual predators (Gonçalves Rodrigues & Silva Absalão, 2005; Mendonça *et al*., 2015; Sigwart, 2018).

Here, we first spectrally characterize the biofluorescence of chitons shells and use visual models to better understand whether the biofluorescence could contribute visually to chromatic and achromatic components of their exitance. Our attention is directed towards the lateral sides of boulders, areas notoriously known to be inhabited by chitons, to test whether the fluorescence present in these patches have the potential of camouflaging the chitons from their predators. In this sense, we characterize the visual system of four potential predators (*i.e.,* octopus, crab, fish, and seagull), identified the reflective and fluorescent components of light that leave chitons’ shells and background substrate (*i.e.,* algae and rocks), and combined these data with field measurements of irradiant spectra (*i.e.*, environmental light), from the chiton’s habitat. Our hypotheses are 1) that fluorescence contributes to enhanced exitance brightness, reducing the achromatic contrast without affecting the chromatic result, and 2) that chitons appear camouflaged to visual predators when on fluorescence backgrounds.

## Material and Methods

### Animal model

The chiton *Ischnoplax pectinata* (G.B. Sowerby II, 1840) is endemic to the West Atlantic, living from a depth of 0-55 m (Gomes, 2015). There is a lack of scientific literature on the species, being restricted to taxonomic studies and compilations of its distribution (Junior, 1985; Oliveira *et al*., 1992; Gomes, 2015). Their occurrence seems to be common on rocky intertidal reefs, where they are found under loose boulders (Pers. Observ.). It seems that their spatial distribution patterns are comparable to those of other chiton species, with the majority of boulders being unoccupied and only a few being occupied (Chapman, 2002; Palmer, 2012).

### Fluorescent body pattern

In October 2021, we collected 44 individuals of *I. pectinata* from an intertidal reef in Northeast Brazil (06°00’22’’S 35°06’24’’W) by searching under boulders. We transported them to the laboratory, along with a boulder sample, and kept them alive in an aerated aquarium for one week. In the laboratory, we measured the size (plate I to VIII), took photos, and collected spectral measurements. Photographs were taken under natural light and in a dark room using a blue light (∼450nm, Fig.S1) to excite fluorescence, with a long pass filter (500 nm). We positioned the camera (Olympus TG-6, ISO 1600, 1/6-13seg, f/3.6 – 4.7) using a tripod and imaged the chitons, including a size scale in each photograph, and saved them in JPG format. Fluorescent areas on chiton’s shells were measured using the package PAT-GEOM in ImageJ (v1.53) (Chan *et al*., 2019). No animal was sacrificed, and all chitons were returned to the sampled site after the procedures.

### Spectral measurements

Three chitons were used for reflectance and fluorescence measurements. Reflectance was measured using an Ocean Optics USB4000-UV-VIS spectrometer, connected to a computer with SpectraSuit (v1.6) software, calibrated with a standard light source (DH-2000-CAL), with a bifurcated fiber optic (QR450-7-XSR), held with a laboratory clamp at 45◦ to normal and positioned 3 mm to the measurement surface (Badiane *et al*., 2017), illumination was provided by a deuterium-tungsten halogen lamp (DH-2000-BAL), integration time set to 51ms, scans to average to 10, and boxcar width to 5. The same set up was used for measured the fluorescence emission spectra, replacing the illumination by the blue light (∼450nm) and including a yellow barrier filter between the chitons and the probe to reduce the excess of excitation light reflected. Any residual excess of excitation light detected by the spectrometer was removed *a posteriori* to avoid misreading of the fluorescence emission spectra (Mazel, 2017). Spectral measurements are restricted to point information on fluorescence patches and reference measurements were made from a Spectralon Reflectance Standard (WS-1-SL) under the respective illumination conditions. Raw spectra were smoothed (*span*=0.1). Additionally, two reflectance measurements were made for crustose red algae backgrounds present on the side of the boulders (*Lithothamnium* sp. and *Peyssonnelia* sp.). For each crustose red algae, spectral data were averaged and smoothed. The resultant spectrum was taken as the background reflectance of both *Lithothamnium* and *Peyssonnelia*. Brightness was calculated as mean relative reflectance in the 300–800 nm interval (Delhey *et al*., 2003; Siefferman & Hill, 2005). We used the same spectrometer with a fiber optic (QP450-2-XSR) and cosine corrected sensor (CC-3-UV) to measure local irradiance. This measurement was taken during low tide (outside of the water) at 09:30h on a cloudless sunny day (Fig.S2). Irradiance measurements (µW. cm^-2^. nm^-1^) were transformed into photon irradiance (photons. s^-1^. cm^-2^. nm^-1^) by multiplication with *h.c* at each wavelength (Fuchs, 2001). We used the practical fluorescent efficiency to calculate the fluorescence contribution of exitance light (Mazel, 2017).

### Visual Modelling

We chose estimates of vision parameters for the visual systems of four potential visually guided predators of chitons: seagulls (Moore, 1975), reef fishes (Randall, 1967), crabs (Silva *et al*., 2010), and octopuses (Mather & Nixon, 1990). Since these predators have not already had all the parameters of their visual systems previously studied, we completed our visual models with available information from species that are phylogenetically close to these predators, or that present similar visual systems. We produced photoreceptor sensitivity curves (Fig.S3) using the available peak sensitivities from Common octopus (*Octopus vulgaris*: 475 nm; Brown & Brown, 1958), European green crab (*Carcinus maenas*: 440, 508 nm; Martin & Mote, 1982), White-banded triggerfish (*Rhinecanthus aculeatus*: 412, 480, 528 nm; Green *et al*., 2022), and the Blue tit (*Cyanistes caeruleus*: 372, 453, 539, 607 nm; Hart *et al*., 2000). We used Blue tits as a proxy to seagulls because both express SWS1 cone pigments that are tuned to the ultraviolet light range, instead of violet (Ödeen & Håstad, 2013). We calculated the absolute photoreceptor quantum catch (*qcatch* = “Qi”) of each predator’s visual system between 300-700nm using the reflectance spectra of the chiton (with and without fluorescence component) and crustose red algae backgrounds (*Lithothamnium* sp. and *Peyssonnelia* sp.), under the local incident light, outside of the water (cloudless sunny day, at 09:30am).

For chromatic contrast models (ΔS), we used the receptor noise limited (RNL) model to predict the of perceptual distance or measurement of contrast discriminability between the chitons and their backgrounds. The RNL model assumes that signal discrimination under optimal condition is constrained by noise originating in the receptors and subsequent neural processing (Vorobyev & Osorio, 1998; Vorobyev *et al*., 2001). Octopuses was excluded for chromatic models as their mechanisms of colour vision have not been conclusively investigated (Hanke & Kelber, 2020). We used a homogeneous transmission of 1 for the ocular media (*trans* = ‘ideal’) for crabs and fish, while for seagulls we employed *trans* = ‘bluetit’ (Hart *et al*., 2000). We applied von Kries transformation to normalize receptor quantum catches to the background, thus accounting for receptor adaptation (*vonkries* = TRUE). The neural photoreceptor noise (*noise* = ‘neural’) was set with a Weber fraction of 0.12 for crabs (Silva *et al*., 2022), 0.05 for fish (Wilkins *et al*., 2016), and 0.1 for birds (Olsson *et al*., 2018). For the relative photoreceptor densities, we employed 1:1 for crabs, since no information about the proportion of photoreceptors is mentioned by literature, 1:2:2 for triggerfish (Champ *et al*., 2014) and 1:1.9:2.7:2.7 for seagulls, based on Blue tit (Hart *et al*., 2000). The remaining parameters were set to default. The chromatic visual models were implemented using the ’*pavo*’ 2.7 R package (Maia *et al*., 2019).

For achromatic contrast models (ΔL), we calculated as proposed by Olsson *et al*. (2018), since its calculations by RNL model warrant caution (van den Berg *et al*., 2020a). Achromatic values were calculated according:

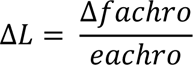

where Δ*f_achro_* is the contrast in achromatic receptor channel and *e_achro_* is the noise parameter. The contrast in achromatic channel was calculated using the spectral sensitivity of the triggerfish double cone (reef fish), the bluetit double cone (seagulls), the longest-wavelength photoreceptor (crabs), and the only one in octopus, as:

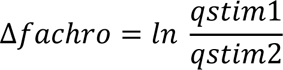

where *q_stim_* is the receptor quantum catch for each stimulus.

To estimate the receptor noise levels (e_achro_) as proposed by Olsson *et al*. (2018) (see also van den Berg *et al*., 2020), we assumed a behavioural determined contrast sensitivity CS of 33.31 to octopus (Nahmad-Rohen & Vorobyev, 2020) and used the Michelson contrast of 0.66 to reef fish (*Rhinecanthus aculeatus*, van den Berg *et al*., 2020). Thus, we obtain the following channel noise estimates: octopus (e_achro_=0.060), reef fish (e_achro_=0.132). To crabs and seagulls, we respectively assumed e_achro_=0.16, based on *Apis mellifera*, and e_achro_=0.34, based on the common starling (see Table 1, Olsson *et al*., 2018), since no information about this is mentioned by literature.

### Statistics

All statistical analyses, colourimetry measurements and visual models were performed in R v4.2.1 (R Core Team, 2020). We tested our data for normality using the Shapiro– Wilk test and homogeneity of variance with Fligner-Killeen test. Total length and fluorescent area on chiton shells were compared by rank using Spearman’s correlations. Mean brightness with/without fluorescence component was tested by bootstrap t-test. Results were considered significant if p < 0.05. The output generated by the chromatic contrast (ΔS) and achromatic contrast (ΔL) are often expressed in units of ‘just-noticeable differences’ (JNDs). We assumed a conservative threshold of 1JND=3ΔS/ΔL (Olsson *et al*., 2018; van den Berg *et al*., 2020a), where < 1JND are considered cryptic, 1-3 JND are poorly discriminable, and > 3 JND are highly discriminable (Siddiqi *et al*., 2004). We interpreted the differences between the fluorescent visual models through visual inspection, comparing the percentile ranges of achromatic and chromatic contrast results between chitons and the background for each visual system.

## Results

The shell plates of *I. pectinata* presented a far-red component (up to and beyond 700 nm) to their reflectance. Under full spectrum irradiance, all reflectance emissions had a primary peak at 800 nm and a secondary peak at roughly 593 nm (Figure 1A). Interestingly, both crustose red algae also reflected longer wavelength (*Peyssonelia*, ∼789 nm; *Lithothamnium*, ∼787 nm), and their reflectance spectra were remarkably similar to each other and to the reflectance of the chiton shells (Figure 1C). When illuminated with higher energy radiation (∼450 nm), we observed that living chitons *I. pectinata* produce a bright “redness” fluorescence to our eyes. This fluorescence was limited to the shell plates. No fluorescence was found on the girdle, but in some individuals, a slight fluorescence was found on the fleshy foot. Two chitons had a similar shape in the fluorescence spectra distribution with maximum emission occurring at a wavelength of 714 nm and a secondary peak at 533 nm (Figure 1B). However, one chiton had a fluorescence emission that was roughly different in shape (primary peak: 757 nm; secondary peak: 591 nm).

**FIGURE 1.**
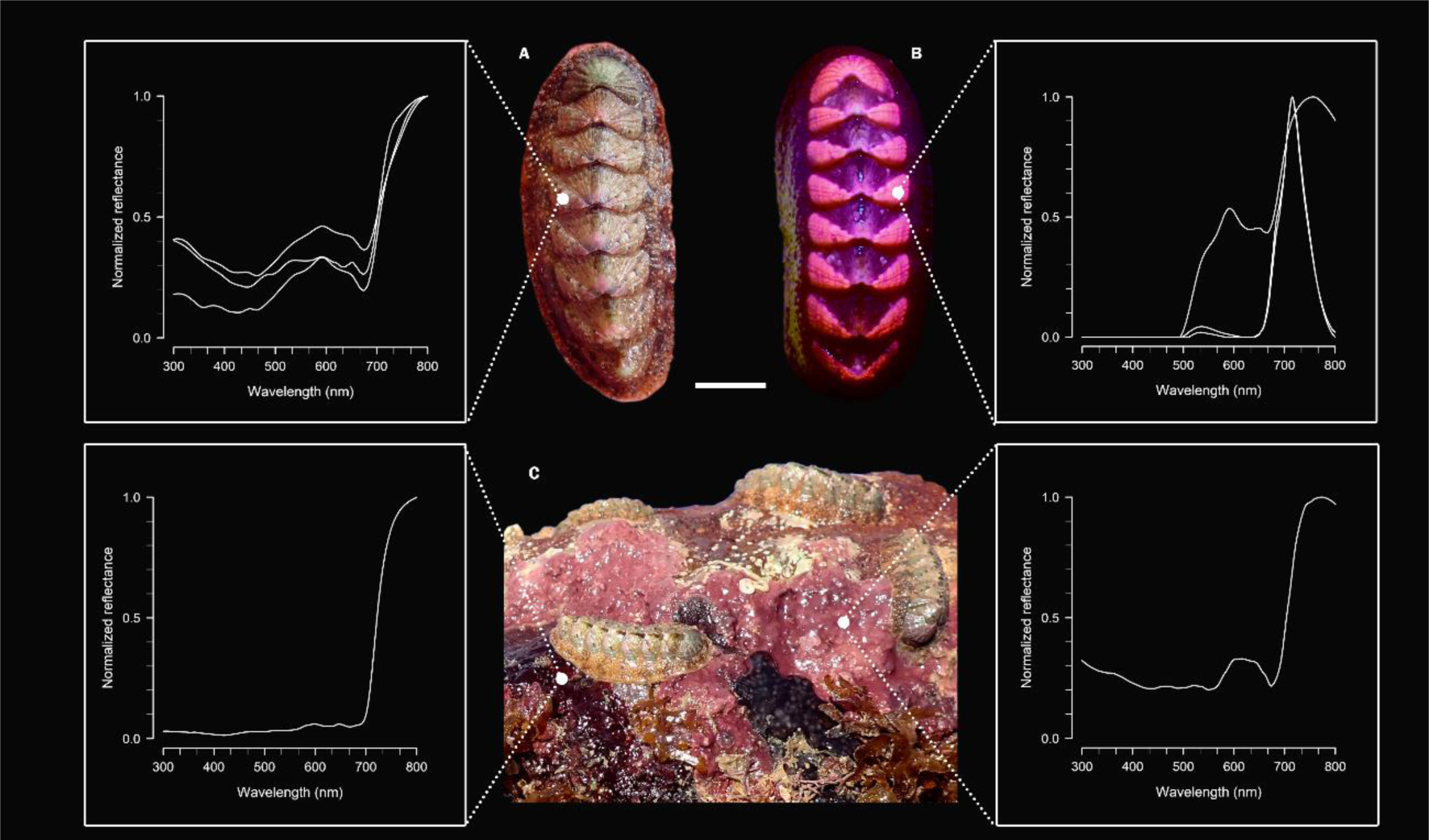
(A). Exitance spectral distribution of *I. pectinata* under full spectrum irradiance, (B) and under fluorescence condition, showing a ‘disruptive’ pattern in shell; (C) Crustose red algae in the boulder they inhabit, *Peyssonnelia* sp. (left) and *Lithothamnium* sp. (right).

Fluorescence was detected in all plates, however in the intermediate valves (II to VII) it appears mainly restricted to the lateral area of the plates producing a fluorescent disruptive pattern. This colour patch pattern shows variability among the 44 individuals. Some individuals displayed fluorescence covering more than 70% of the shell surface area, while others had less than 10% coverage (Figure 2). A weak negative correlation was observed between the total length (mean: 3.04±0.43) and the fluorescence area (mean: 0.74±0.45) of chiton shells (rho = -0.36, p = 0.017).

**FIGURE 2.**
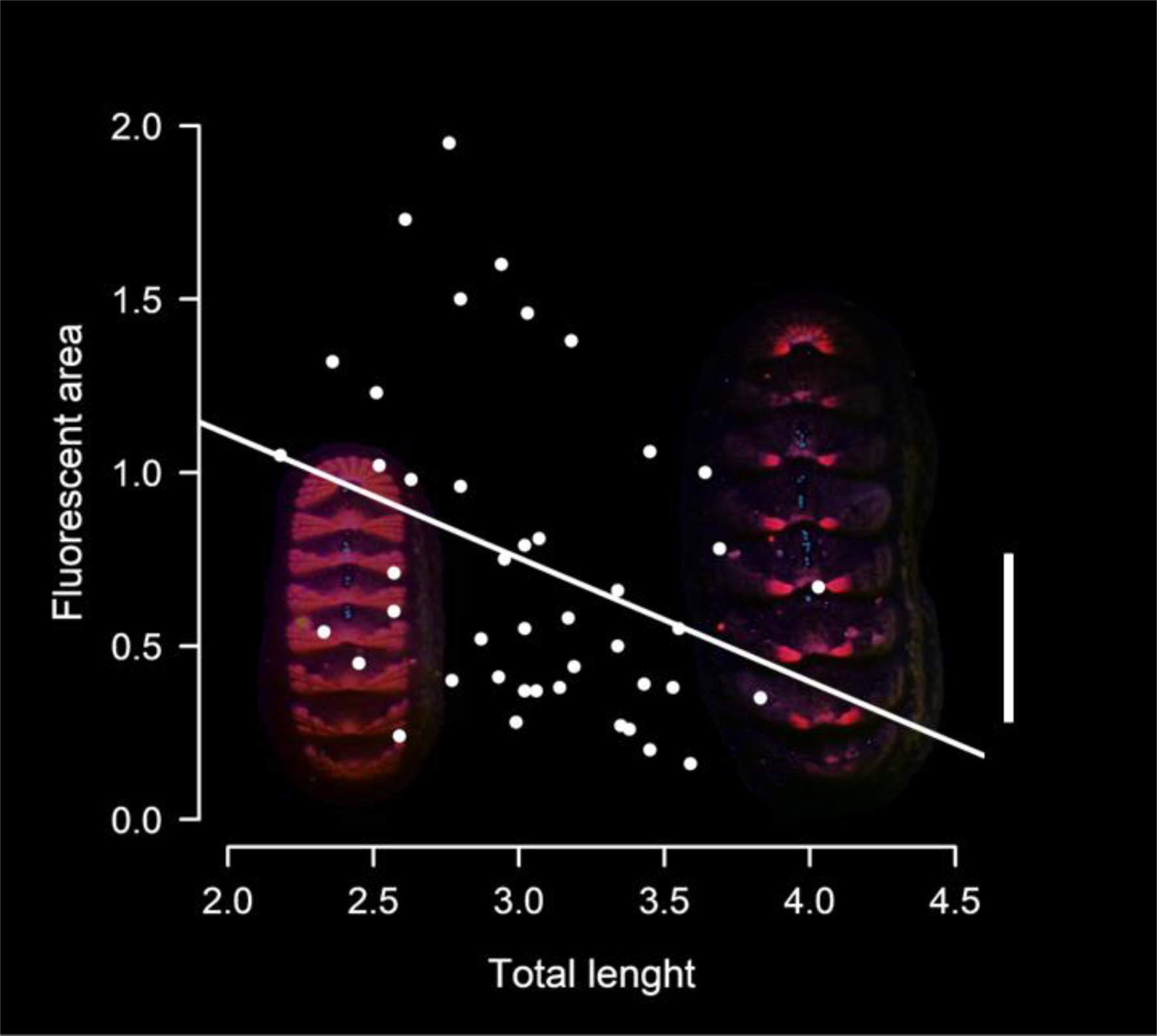
Relationship between total length (cm) and total of fluorescent area (cm^2^) on the shell of *I. pectinata*. Line, fit a linear regression with negative correlation illustrating the rho = -0.36, p = 0.017. Wallpaper, a specimen with 78% fluorescence area (left) and other with less of 8% (right). Scale Bar = 1 cm.

Given our results, the quantified fluorescence contributes, on average, to 21.11% of the exitance light between 650-800 nm, considering the environmental light spectrum (*i.e.*, illuminant) of chitons’ natural habitat (at 09:30h). While, for two of the chitons, this contribution only represented an average of 10%, for a third chiton, it reached an average of 39% (Fig. S4). When we compared the exitance light including fluorescence (mean: 6.48±1.95) to that without fluorescence (mean: 0.36±0.13, bootstrapped T test, p-value = 0.04), we found that the fluorescence enhances the mean brightness of individuals.

### Visual Models

The presence of fluorescence in chitons’ exitance leads to a reduction in discrimination values (JND), especially for achromatic contrast. This reduction is context-dependent, with more pronounced effect on *Lithothamnium* than *Peyssonnelia* background (Figure 3). For all visual systems, fluorescence considerably reduces the achromatic discrimination, except for octopuses, where the presence of fluorescence appears to have little effect. For others, this reduction is sufficient to make chitons poorly discriminated in most case. Exception was a decrease from high to marginally high discriminability for fish when over *Lithothamnium* backgrounds and a shift from poor to marginally cryptic discriminability for gull on *Peyssonnelia* backgrounds. In terms of chromatic effects, fluorescence does not appear to influence colour visual perception. While in specific visual contexts, fluorescence may reduce discrimination levels (*e.g.,* fish, on *Peyssonnelia*; gull, on *Lythothamnium*), visual confirmation is challenging due to overlapping interquartile ranges (Figure 3).

**FIGURE 3.**
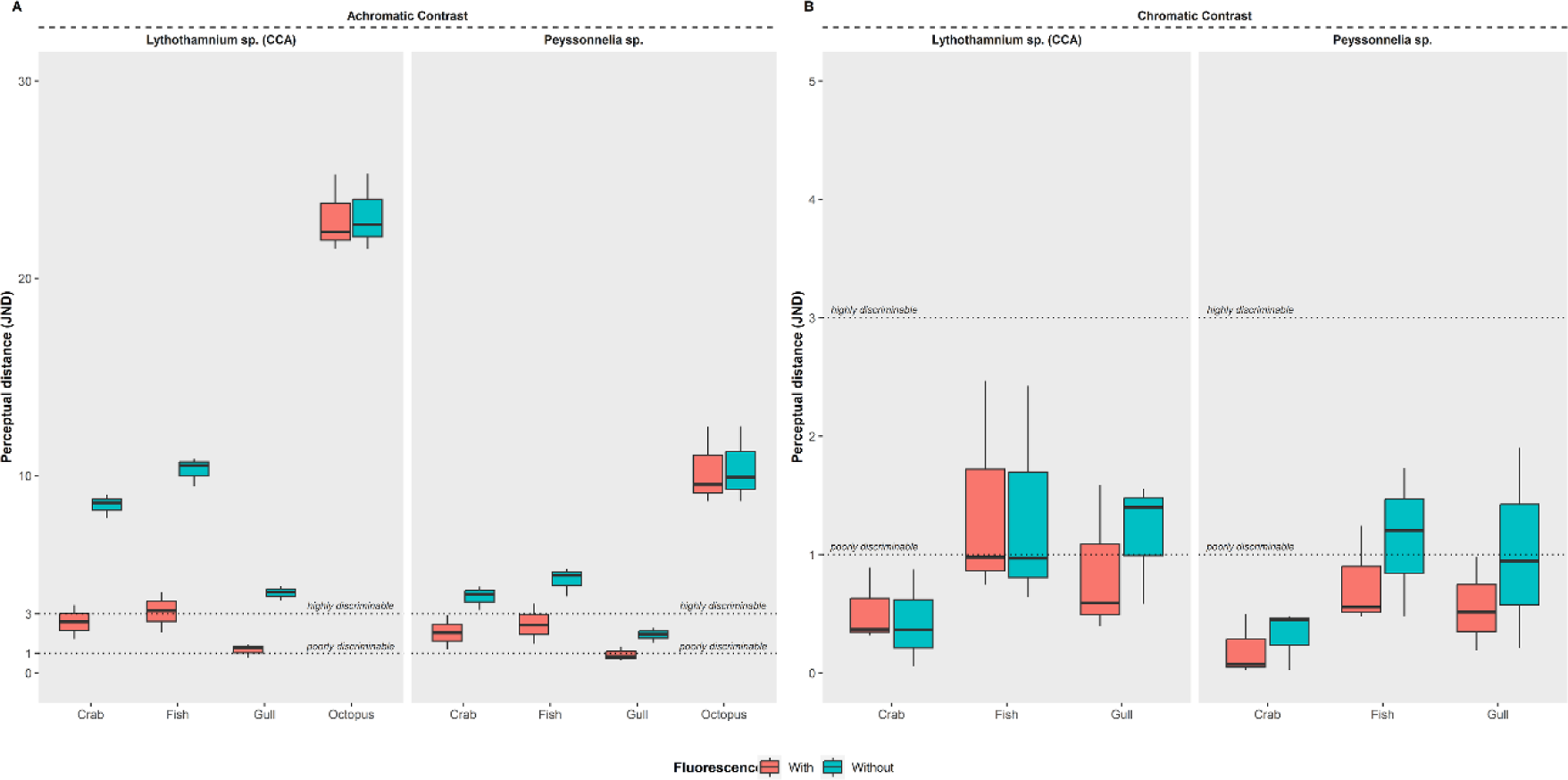
A) Achromatic contrast (ΔL) and (B) Chromatic contrast (ΔS) between the natural exitance spectra (with and without fluorescence) of chiton shells and the reflectance spectra of two crustose red algae (*Lithothamnium* sp. and *Peyssonnelia* sp.), modelled based on the visual systems of different predators.

## Discussion

The first description of biofluorescence in chitons (*Ischnoplax pectinate*), an ancient group of Mollusca, reinforces the relevance of traits influenced by vision-related biological drivers across different groups and moments of evolutionary history. To the best of our knowledge, biofluorescence in molluscs has only been reported in nudibranchs (Betti *et al*., 2021), gastropods (Williams *et al*., 2016), and bivalves (Wanamaker *et al*., 2009), but not yet in polyplacophorans. Our findings provide evidence for the visual function of biofluorescence, although its effect on concealment of chitons is context-dependent according to background, chromatic and achromatic aspects of vision, and type of predator.

Fluorescence has been proposed to have a signalling function in mantis shrimps (Mazel, 2004), jumping spiders (Lim *et al*., 2007), fish (Gerlach *et al*., 2014), and budgerigars (Arnold, 2002). Behavioural experiments demonstrate that fairy wrasses perceive and respond to their red fluorescent colouration in a mirror image stimuli experiment (Gerlach *et al*., 2014). Our evidence supports the hypothesis that fluorescence may have a visual role. Our results demonstrate that it contributes to an increase in the brightness of chiton shells, affecting the achromatic contrast discrimination in the vision of the most of their visual predators, even in daylight conditions. This is accomplished as fluorescence adds photons to the long-wavelength range through the conversion of irradiance from the blue portion of the spectrum (Johnsen, 2011). While our results showed a pronounced effect of biofluorescence in reducing achromatic contrast, its effects on chromatic contrast remained uncertain. Nevertheless, altering the spectral distribution shape of exitance light has the potential to impact colour appearance of objects (Shevell, 2003). Undoubtedly, the contribution of the fluorescent component can vary depending on the ambient light conditions (Bitton *et al*., 2017; Taboada *et al*., 2017). In their study, Taboada et al. (2017) discovered that fluorescence exhibited a 1.6 times higher contribution when calculated using the irradiance of the full moon compared to twilight, which has a higher relative proportion of ‘blue’ radiation in its composition (Cronin *et al*., 2014). We expect fluorescence to become especially relevant to the exitance light under spectrally skewed environments (*e.g.,* increasing depth), reinforcing its visual function. It is interesting to note that biofluorescence in *I. pectinata* contributes particularly to the 650-800 nm spectral range, while the reflectance of both chitons and crustose red algae also falls within the red, or far-red, range (up to and beyond 700 nm), outside the optical sensitivity range of most animal visual systems (approximately 300-700 nm). This could provide effective camouflage in the bluer depths, in which chitons can inhabit (∼50 m), similar to the effect that gives a bluish appearance to fishes that have red, or far-red components in their reflectance (Marshall & Johnsen, 2011). Variation in ambient light with depth may exchange the functional role of fluorescence with other potential roles, such as photoprotection (Salih *et al*., 2000) and warning signals (Nemésio, 2005). Furthermore, we also expect fluorescence to be accentuated in the algae’s background, as they are also fluorescent, although our study has not determined their fluorescent emission spectrum.

Our data shows that as individuals become larger, the surface area of fluorescence of their shells becomes smaller. This suggests a cost–benefit trade-off that requires allocation of limited resources at the expense of allocation to other fitness components, such as growth, survivorship, and fecundity, which consequently can induce variation in camouflage strategies throughout ontogeny. Our study did not include samples from young individuals, but field records show that, at this stage, fluorescence covers almost the entire shell surface (unpublished results, Fig. S5). A suitable approach to investigate whether the colour patterns of *I. pectinata* can function as different camouflage strategies throughout ontogeny, with juvenile adopting a ‘background matching’ strategy and adults a ‘disruptive colouration’, is to use photography to visual modelling how these patterns might look through the eyes of their predators by a analytical framework, as QCPA (van den Berg *et al*., 2020b) or MICA Toolbox (Troscianko & Stevens, 2015; Troscianko *et al*., 2017).

It is likely that the biofluorescence in *I. pectinata* cannot be synthesized by itself but must be acquired from dietary sources as fluorescent compounds, that can be obtained from crustose red algae, with subsequent biomineralization in the shell.

Notwithstanding, crustose coralline algae (CCA) represent the major food item found in the gut content of many chitons species (Demopulos, 1975; Nishi, 1975; Roob, 1975; Piercy, 1987; Sigwart & Schwabe, 2017) and it may be an important factor in their spatial distribution (Kangas & Shepherd, 1984; Reyes-Gómez *et al*., 2010; Liversage, 2016). CCA are especially important as members of coral reef communities because they are builders of carbonate material and contribute to the biotic spatial structure of reefs (Littler & Littler, 1984), facilitating the larval settlement of several groups of benthic invertebrates (Gee, 1965; Barnes & Gonor, 1973; Price, 2010). This chiton-coralline reef-building association has been acknowledged in few studies, but its importance for benthic communities structuring has been neglected (Littler *et al*., 1995).

Previous studies have hypothesized that chiton’s shell colours could be driven by their visual guide predators acting as selective agents (Gonçalves Rodrigues & Silva Absalão, 2005; Mendonça *et al*., 2015; Sigwart, 2018). Experimentally, a behavioural study showed that chitons that were colour camouflaged to their substrate, according to human vision, were still frequently attacked by shore crabs (Mendonça *et al*., 2016). Our visual models suggest that the chiton-crustose red algae substratum camouflage must be chromatically effective while also being poorly discriminable in achromatic terms to the crab’s vision, indicating that these interactions could be mediated by achromatic aspects of vison, as well as mechanical and/or chemosensory cues.

## Conclusion

Here, we applied an over-simplified set of assumptions for predator visual systems. It’s important to note that deviations from these model assumptions could potentially impact the overall conclusions presented. Therefore, additional physiological studies and behavioural threshold experiments on the visual systems of chiton’s predators is necessary to refine our visual models. Additionally, evaluating different backgrounds, lightning conditions, and other factors that may influence chromatic and achromatic perception will contribute to a more comprehensive understanding (Sibeaux *et al*., 2019; van den Berg *et al*., 2020a; Venables *et al*., 2022). Furthermore, it is unknown whether fluorescence intensity changes over an individual’s lifetime or whether dietary and seasonal variations can affect it. Molecular studies to trace the chemical origin of fluorescence in *I. pectinata* are therefore warranted. Our results provide evidence for our hypothesis that fluorescence contributes to enhanced exitance brightness, leading to a reduction in achromatic contrast without influencing the chromatic outcome. Nevertheless, the context-dependent factor on chitons’ camouflage requires further investigation through behavioural experiments. Our findings demonstrate that biofluorescence can affect the visual aspects of ecological interactions, increasing the brightness and influencing the visual perception of predators.

## Supporting information

File S2_Colour patch data

## Acknowledgements

We would like to express our gratitude to Sofia Coradini, Diogo Jackson, and Marilia Erickson for their initial laboratory support. Special thanks go to Noelle Lechat, Colonia de Pesca Z-10 (Miguel and Susana), and Oceânica – Pesquisa, Educação e Conservação for providing the necessary logistical support during the fieldwork. Additionally, we extend our heartfelt appreciation to Dr. Mazel for his encouraging guidance. Finally, we would like to thank the editor and reviewers for their valuable comments.

## Supporting Information

Additional supporting information may be found in the online version of this article on the publisher’s website.

File S1. Supplementary figures.docx

File S2. Colour patch data.csv

## Data Availability

The data underlying this article are available in Zenodo, at 10.5281/zenodo.8279507. Other data are available in the Supporting Information.

## Author Contributions

Conceptualization: G.G.G., R.S.G., D.M.A.P, and P.A.H.

Data curation: G.G.G

Formal Analysis: G.G.G, and M.H.

Investigation: G.G.G.

Methodology: G.G.G, R.S.G., D.M.A.P, and P.A.H

Project administration: G.G.G.

Resources: G.G.G., D.M.A.P, and P.A.H

Software: G.G.G.

Supervision: D.M.A.P, and P.A.H.

Visualization: G.G.G.

Writing – original draft: G.G.G., R.S.G., J.A.J., and P.A.H.

Writing – review & editing: G.G.G, R.S.G., J.A.J., M.H., D.M.A.P, and P.A.H.

## Ethics

The field location is publicly accessible, and no further licenses were needed. This study was exempt by the Ethics Committee on the Use of Animals, following Brazilian federal legislation (n°11.794, October 8, 2008).

## References

Aihara Y, Maruyama S, Baird AH, Iguchi A, Takahashi S, Minagawa J. 2019. Green fluorescence from cnidarian hosts attracts symbiotic algae. Proceedings of the National Academy of Sciences 116: 2118–2123.

Arnold KE. 2002. Fluorescent Signaling in Parrots. Science 295: 92–92.

Badiane A, Pérez i de Lanuza G, Font E. 2017. Colour patch size and measurement error using reflectance spectrophotometry. Methods in Ecology and Evolution 8: 1585– 1593.

Barnes JR, Gonor JJ. 1973. The larval settling response of the lined chiton Tonicella lineata. Marine Biology 20: 259–264.

van den Berg CP, Hollenkamp M, Mitchell LJ, Watson EJ, Green NF, Marshall NJ, Cheney KL. 2020a. More than noise: Context-dependant luminance contrast discrimination in a coral reef fish (*Rhinecanthus aculeatus*). Journal of Experimental Biology: jeb.232090.

van den Berg CP, Troscianko J, Endler JA, Marshall NJ, Cheney KL. 2020b. Quantitative Colour Pattern Analysis (QCPA): A comprehensive framework for the analysis of colour patterns in nature (G Iossa, Ed.). Methods in Ecology and Evolution 11: 316–332.

Betti F, Bavestrello G, Cattaneo-Vietti R. 2021. Preliminary evidence of fluorescence in Mediterranean heterobranchs. Journal of Molluscan Studies 87: eyaa040.

Bitton PP, Harant UK, Fritsch R, Champ CM, Temple SE, Michiels NK. 2017. Red fluorescence of the triplefin *Tripterygion delaisi* is increasingly visible against background light with increasing depth. Royal Society Open Science 4: 161009.

Bracken MES, Oates JM, Badten AJ, Bernatchez G. 2018. Predicting rates of consumer-mediated nutrient cycling by a diverse herbivore assemblage. Marine Biology 165: 165.

Brown PK, Brown PS. 1958. Visual Pigments of the Octopus and Cuttlefish. Nature 182: 1288–1290.

Caro T, Argueta Y, Briolat ES, Bruggink J, Kasprowsky M, Lake J, Mitchell MJ, Richardson S, How M. 2019. Benefits of zebra stripes: Behaviour of tabanid flies around zebras and horses. PLoS ONE 14: e0210831.

Caro T, Stoddard MC, Stuart-Fox D. 2017. Animal coloration research: why it matters. Philosophical Transactions of the Royal Society B: Biological Sciences 372: 20160333.

Champ C, Wallis G, Vorobyev M, Siebeck U, Marshall J. 2014. Visual Acuity in a Species of Coral Reef Fish: Rhinecanthus aculeatus. Brain, Behavior and Evolution 83: 31–42.

Chan IZW, Stevens M, Todd PA. 2019. pat-geom: A software package for the analysis of animal patterns (D Silvestro, Ed.). Methods in Ecology and Evolution 10: 591–600.

Chapman MG. 2002. Patterns of spatial and temporal variation of macrofauna under boulders in a sheltered boulder field. Austral Ecology 27: 211–228.

Chappell DR, Speiser DI. 2023. Polarization sensitivity and decentralized visual processing in an animal with a distributed visual system. Journal of Experimental Biology 226: jeb244710.

Connors MJ, Ehrlich H, Hog M, Godeffroy C, Araya S, Kallai I, Gazit D, Boyce M, Ortiz C. 2012. Three-dimensional structure of the shell plate assembly of the chiton Tonicella marmorea and its biomechanical consequences. Journal of Structural Biology 177: 314–328.

Cronin TW, Johnsen S, Marshall NJ, Warrant EJ. 2014. Visual Ecology. New Jersey: Princeton University Press.

Davis Rabosky AR, Cox CL, Rabosky DL, Title PO, Holmes IA, Feldman A, McGuire JA. 2016. Coral snakes predict the evolution of mimicry across New World snakes. Nature Communications 7: 11484.

Delhey K, Johnsen A, Peters A, Andersson S, Kempenaers B. 2003. Paternity analysis reveals opposing selection pressures on crown coloration in the blue tit (Parus caeruleus). Proceedings of the Royal Society of London. Series B: Biological Sciences 270: 2057–2063.

Demopulos PA. 1975. Diet, Activity and Feeding in Tonicella lineata (Wood,1815). The Veliger 18: 42–46.

Endler JA. 1978. A Predator’s View of Animal Color Patterns. In: Hecht MK., In: Steere WC., In: Wallace B, eds. Evolutionary Biology. Evolutionary Biology. Boston, MA: Springer US, 319–364.

Endler JA. 1993. The Color of Light in Forests and Its Implications. Ecological Monographs 63: 1–27.

Endler JA, Houde AE. 1995. GEOGRAPHIC VARIATION IN FEMALE PREFERENCES FOR MALE TRAITS IN POECILIA RETICULATA. Evolution 49: 456–468.

Eyal G, Wiedenmann J, Grinblat M, D’Angelo C, Kramarsky-Winter E, Treibitz T, Ben-Zvi O, Shaked Y, Smith TB, Harii S, Denis V, Noyes T, Tamir R, Loya Y. 2015. Spectral Diversity and Regulation of Coral Fluorescence in a Mesophotic Reef Habitat in the Red Sea (CR Voolstra, Ed.). PLOS ONE 10: e0128697.

Fernandez CZ, Vendrasco MJ, Runnegar B. 2007. Aesthete Canal Morphology in Twelve Species of Chiton (Polyplacophora). The Veliger 49: 51–69.

Fuchs E. 2001. Separating the fluorescence and reflectance components of coral spectra. Applied Optics 40: 3614.

Gagnon P, Matheson K, Stapleton M. 2012. Variation in rhodolith morphology and biogenic potential of newly discovered rhodolith beds in Newfoundland and Labrador (Canada). Botanica Marina 55: 85–99.

Gandía-Herrero F, García-Carmona F, Escribano J. 2005. Floral fluorescence effect. Nature 437: 334–334.

Gee JM. 1965. Chemical stimulation of settlement in larvae of Spirorbis rupestris (Serpulidae). Animal Behaviour 13: 181–186.

Gerlach T, Sprenger D, Michiels NK. 2014. Fairy wrasses perceive and respond to their deep red fluorescent coloration. Proceedings of the Royal Society B: Biological Sciences 281: 20140787.

Gomes JAJ. 2015. Revisão taxonômica do gênero Ischnoplax (Chitonoidea; Callistoplacidae) do Atlântico Oeste. Unpublished thesis, Universidade de São Paulo.

Gonçalves Rodrigues LR, Silva Absalão R. 2005. Shell colour polymorphism in the chiton Ischnochiton striolatus (Gray, 1828) (Mollusca: Polyplacophora) and habitat heterogeneity. Biological Journal of the Linnean Society 85: 543–548.

Green NF, Guevara E, Osorio DC, Endler JA, Marshall NJ, Vorobyev M, Cheney KL. 2022. Colour discrimination thresholds vary throughout colour space in a reef fish (*Rhinecanthus aculeatus*). Journal of Experimental Biology 225: jeb243533.

Haddock SHD. 2005. Bioluminescent and Red-Fluorescent Lures in a Deep-Sea Siphonophore. Science 309: 263–263.

Hagen JFD, Roberts NS, Johnston RJ. 2023. The evolutionary history and spectral tuning of vertebrate visual opsins. Developmental Biology 493: 40–66.

Hanke FD, Kelber A. 2020. The Eye of the Common Octopus (Octopus vulgaris). Frontiers in Physiology 10: 1637.

Hart NS, Partridge JC, Cuthill IC, Bennett AT. 2000. Visual pigments, oil droplets, ocular media and cone photoreceptor distribution in two species of passerine bird: the blue tit (Parus caeruleus L.) and the blackbird (Turdus merula L.). Journal of Comparative Physiology. A, Sensory, Neural, and Behavioral Physiology 186: 375–387.

Heiling AM, Herberstein ME, Chittka L. 2003. Crab-spiders manipulate flower signals. Nature 421: 334–334.

Johnsen S. 2011. The Optics of Life: A Biologist’s Guide to Light in Nature. Princeton, NJ: Princeton Univ Pr.

Junior OG. 1985. Sobre Ischnoplax pectinatus (Sowerby, 1840) e sua ocorrência no litoral sul do Brasil (Mollusca, Polyplacophora). Memórias do Instituto Oswaldo Cruz 80: 401–406.

Kangas M, Shepherd SA. 1984. Distribution and feeding of chitons in a boulder habitat at West Island, South Australia. Journal of the Malacological Society of Australia 6: 101–111.

Kelber A, Osorio D. 2010. From spectral information to animal colour vision: experiments and concepts. Proceedings of the Royal Society B: Biological Sciences 277: 1617–1625.

Kelber A, Vorobyev M, Osorio D. 2003. Animal colour vision – behavioural tests and physiological concepts. Biological Reviews of the Cambridge Philosophical Society 78: 81–118.

van der Kooi CJ, Stavenga DG, Arikawa K, Belušič G, Kelber A. 2021. Evolution of Insect Color Vision: From Spectral Sensitivity to Visual Ecology. Annual Review of Entomology 66: 435–461.

Lahti DC, Ardia DR. 2016. Shedding Light on Bird Egg Color: Pigment as Parasol and the Dark Car Effect. The American Naturalist 187: 547–563.

Lamb JY, Davis MP. 2020. Salamanders and other amphibians are aglow with biofluorescence. Scientific Reports 10: 2821.

Lim AYH, Chan IZW, Carrasco LR, Todd PA. 2019. Aposematism in pink warty sea cucumbers: independent effects of chromatic and achromatic cues. Marine Ecology Progress Series 631: 157–164.

Lim MLM, Land MF, Li D. 2007. Sex-Specific UV and Fluorescence Signals in Jumping Spiders. Science 315: 481–481.

Littler MM, Littler DS. 1984. Models of tropical reef biogenesis: the contribution of algae. Progress in Phycological Research 3: 323–364.

Littler MM, Littler DS, Taylor PR. 1995. Selective Herbivore Increases Biomass of Its Prey: A Chiton-Coralline Reef-Building Association. Ecology 76: 1666–1681.

Liu C, Liu H, Huang J, Ji X. 2022. Optimized Sensory Units Integrated in the Chiton Shell. Marine Biotechnology 24: 380–392.

Liversage K. 2016. The influence of boulder shape on the spatial distribution of crustose coralline algae (Corallinales, Rhodophyta). Marine Ecology 37: 459–462.

Liversage K, Benkendorff K. 2017. The first observations of *Ischnochiton* (Mollusca, Polyplacophora) movement behaviour, with comparison between habitats differing in complexity. PeerJ 5: e4180.

Maia R, Gruson H, Endler JA, White TE. 2019. pavo 2: New tools for the spectral and spatial analysis of colour in r (RB O’Hara, Ed.). Methods in Ecology and Evolution 10: 1097–1107.

Marshall J, Johnsen S. 2011. Camouflage in marine fish. In: Stevens M, In: Merilaita S, eds. Animal Camouflage: Mechanisms and Function. Cambridge: Cambridge University Press, 186–211.

Marshall J, Johnsen S. 2017. Fluorescence as a means of colour signal enhancement. Philosophical Transactions of the Royal Society B: Biological Sciences 372: 20160335.

Marshall KLA, Stevens M. 2014. Wall lizards display conspicuous signals to conspecifics and reduce detection by avian predators. Behavioral Ecology 25: 1325– 1337.

Martin FG, Mote MI. 1982. Color receptors in marine crustaceans: a second spectral class of retinular cell in the compound eyes of Callinectes and Carcinus. Journal of Comparative Physiology 145: 549–554.

Mather J, Nixon M. 1990. Octopus vulgaris drills Chiton. Journal of Cephalopod Biology 1: 113–116.

Mazel CH. 2004. Fluorescent Enhancement of Signaling in a Mantis Shrimp. Science 303: 51–51.

Mazel C. 2017. Method for Determining the Contribution of Fluorescence to an Optical Signature, with Implications for Postulating a Visual Function. Frontiers in Marine Science 4: 266.

Mendonça V, Vinagre C, Boaventura D, Cabral H, Silva ACF. 2016. Chitons’ apparent camouflage does not reduce predation by green crabs *Carcinus maenas*. Marine Biology Research 12: 125–132.

Mendonça V, Vinagre C, Cabral H, Silva ACF. 2015. Habitat use of inter-tidal chitons - role of colour polymorphism. Marine Ecology 36: 1098–1106.

Mercegue V, Ibáñez CM, Sepúlveda RD. 2021. Intertidal microhabitats as a shelter for assemblages of chitons at southern Chile. Regional Studies in Marine Science 46: 101886.

Montecinos C, Riera R, Brante A. 2020. Site fidelity and homing behaviour in the intertidal species Chiton granosus (Polyplacophora) (Frembly 1889). Journal of Sea Research 164: 101932.

Moore M. 1975. Foraging of the western gull, Larus occidentalis and its impact on the chiton Nuttalina californica. The Veliger 18: 51–53.

Nahmad-Rohen L, Vorobyev M. 2020. Spatial Contrast Sensitivity to Polarization and Luminance in Octopus. Frontiers in Physiology 11: 379.

Nemésio A. 2005. Fluorescent colors in orchid bees (Hymenoptera: Apidae). Neotropical Entomology 34: 933–936.

Nilsson DE, Smolka J, Bok M. 2022. The vertical light-gradient and its potential impact on animal distribution and behavior. Frontiers in Ecology and Evolution 10: 951328.

Nishi R. 1975. The Diet and Feeding habits of Nuttallina californica (Reeve, 1847) from two contrasting habitats in Central California. The Veliger 18: 30–33.

Ödeen A, Håstad O. 2013. The phylogenetic distribution of ultraviolet sensitivity in birds. BMC Evolutionary Biology 13: 36.

Oliveira MM, Dijck MPM, Mello RLS. 1992. Polyplacophora (Mollusca) do Nordeste do Brasil. Caderno Omega Universidade Federal Rural de Pernambuco 3: 59–65.

Olson ER, Carlson MR, Ramanujam VMS, Sears L, Anthony SE, Anich PS, Ramon L, Hulstrand A, Jurewicz M, Gunnelson AS, Kohler AM, Martin JG. 2021. Vivid biofluorescence discovered in the nocturnal Springhare (Pedetidae). Scientific Reports 11: 4125.

Olsson P, Lind O, Kelber A. 2018. Chromatic and achromatic vision: parameter choice and limitations for reliable model predictions. Behavioral Ecology 29: 273–282.

Osorio D, Vorobyev M. 1997. Colour vision as an adaptation to frugivory in primates. Proceedings of the Royal Society B: Biological Sciences 263: 593–599.

Palmer ANS. 2012. Spatial and genetic investigation of aggregation in Ischnochiton (Polyplacophora; Neoloricata; Ischnochitonina; Ischnochitonidae; Ischnochitoninae) species with different larval development. Austral Ecology 37: 110–124.

Piercy RD. 1987. Habitat and Food Preferences in Six Eastern Pacific Chiton Species (Mollusca: Polyplacophora). The Veliger 29: 388–393.

Price N. 2010. Habitat selection, facilitation, and biotic settlement cues affect distribution and performance of coral recruits in French Polynesia. Oecologia 163: 747– 758.

Price N, Green S, Troscianko J, Tregenza T, Stevens M. 2019. Background matching and disruptive coloration as habitat-specific strategies for camouflage. Scientific Reports 9: 7840.

Prötzel D, Heß M, Schwager M, Glaw F, Scherz MD. 2021. Neon-green fluorescence in the desert gecko Pachydactylus rangei caused by iridophores. Scientific Reports 11: 297.

R Core Team. 2020. R: A language and environment for statistical computing.

Randall JE. 1967. Food habits of reef fishes of the West Indies. Studies in Tropical Oceanography 5: 665–847.

Renoult JP, Kelber A, Schaefer HM. 2017. Colour spaces in ecology and evolutionary biology: Colour spaces in ecology and evolutionary biology. Biological Reviews 92: 292–315.

Reyes-Gómez A, Barrientos-Luján NA, Medina-Bautista J, Ramírez-Luna S. 2010. Chitons from the coralline area of Oaxaca, Mexico (Polyplacophora). Bollettino Malacologico: 110–124.

Roob MF. 1975. The Diet of the Chiton Cyanoplax hartwegii in three intertidal habitats. The Veliger 18: 34–37.

Saito H, Fujikura K, Tsuchida S. 2008. Chitons (Mollusca: Polyplacophora) associated with hydrothermal vents and methane seeps around Japan, with descriptions of three new species*. American Malacological Bulletin 25: 113–124.

Salih A, Larkum A, Cox G, Hoegh-Guldberg O. 2000. Fluorescent pigments in corals are photoprotective. Nature 408: 850–853.

Shevell SK (Ed.). 2003. The Science of Color. Amsterdam ; Boston : [United States]: Elsevier Science Ltd.

Sibeaux A, Cole GL, Endler JA. 2019. Success of the receptor noise model in predicting colour discrimination in guppies depends upon the colours tested. Vision Research 159: 86–95.

Siddiqi A, Cronin TW, Loew ER, Vorobyev M, Summers K. 2004. Interspecific and intraspecific views of color signals in the strawberry poison frog Dendrobates pumilio. The Journal of Experimental Biology 207: 2471–2485.

Siebeck UE, Parker AN, Sprenger D, Mäthger LM, Wallis G. 2010. A Species of Reef Fish that Uses Ultraviolet Patterns for Covert Face Recognition. Current Biology 20: 407–410.

Siefferman L, Hill GE. 2005. UV-blue structural coloration and competition for nestboxes in male eastern bluebirds. Animal Behaviour 69: 67–72.

Sigwart JD. 2017. Deep trees: Woodfall biodiversity dynamics in present and past oceans. Deep Sea Research Part II: Topical Studies in Oceanography 137: 282–287.

Sigwart JD. 2018. The chiton stripe tease. Marine Biodiversity 48: 1277–1278.

Sigwart JD, Schwabe E. 2017. Anatomy of the many feeding types in polyplacophoran molluscs. Invertebrate Zoology 14: 205–216.

Silva DJA, Erickson MF, dos Santos Guidi R, Pessoa DMA. 2022. Thin-fingered fiddler crabs display a natural preference for UV light cues but show no sensory bias to other hypertrophied claw coloration. Behavioural Processes 200: 104667.

Silva AC, Silva IC, Hawkins SJ, Boaventura DM, Thompson RC. 2010. Cheliped morphological variation of the intertidal crab Eriphia verrucosa across shores of differing exposure to wave action. Journal of Experimental Marine Biology and Ecology 391: 84–91.

Sparks JS, Schelly RC, Smith WL, Davis MP, Tchernov D, Pieribone VA, Gruber DF. 2014. The Covert World of Fish Biofluorescence: A Phylogenetically Widespread and Phenotypically Variable Phenomenon (D Fontaneto, Ed.). PLoS ONE 9: e83259.

Stuart-Fox D, Newton E, Clusella-Trullas S. 2017. Thermal consequences of colour and near-infrared reflectance. Philosophical Transactions of the Royal Society B: Biological Sciences 372: 20160345.

Sumner-Rooney L, Sigwart JD. 2018. Do chitons have a brain? New evidence for diversity and complexity in the polyplacophoran central nervous system. Journal of Morphology 279: 936–949.

Taboada C, Brunetti AE, Pedron FN, Carnevale Neto F, Estrin DA, Bari SE, Chemes LB, Peporine Lopes N, Lagorio MG, Faivovich J. 2017. Naturally occurring fluorescence in frogs. Proceedings of the National Academy of Sciences 114: 3672– 3677.

Thorp RW, Briggs DL, Estes JR, Erickson EH. 1975. Nectar Fluorescence under Ultraviolet Irradiation. Science 189: 476–478.

Troscianko J, Skelhorn J, Stevens M. 2017. Quantifying camouflage: how to predict detectability from appearance. BMC Evolutionary Biology 17: 7.

Troscianko J, Stevens M. 2015. Image calibration and analysis toolbox – a free software suite for objectively measuring reflectance, colour and pattern (S Rands, Ed.). Methods in Ecology and Evolution 6: 1320–1331.

Venables SV, Drerup C, Powell SB, Marshall NJ, Herbert-Read JE, How MJ. 2022. Polarization vision mitigates visual noise from flickering light underwater. Science Advances 8: eabq2770.

Vorobyev M, Brandt R, Peitsch D, Laughlin SB, Menzel R. 2001. Colour thresholds and receptor noise: behaviour and physiology compared. Vision Research 41: 639–653.

Vorobyev M, Osorio D. 1998. Receptor noise as a determinant of colour thresholds. Proceedings of the Royal Society of London. Series B: Biological Sciences 265: 351– 358.

Wanamaker AD, Baker A, Butler PG, Richardson CA, Scourse JD, Ridgway I, Reynolds DJ. 2009. A novel method for imaging internal growth patterns in marine mollusks: A fluorescence case study on the aragonitic shell of the marine bivalve *Arctica islandica* (Linnaeus): Imaging mollusk shells via fluorescence. Limnology and Oceanography: Methods 7: 673–681.

Wilkins L, Marshall NJ, Johnsen S, Osorio D. 2016. Modelling fish colour constancy, and the implications for vision and signalling in water. Journal of Experimental Biology: jeb.139147.

Williams ST, Ito S, Wakamatsu K, Goral T, Edwards NP, Wogelius RA, Henkel T, de Oliveira LFC, Maia LF, Strekopytov S, Jeffries T, Speiser DI, Marsden JT. 2016. Identification of Shell Colour Pigments in Marine Snails Clanculus pharaonius and C. margaritarius (Trochoidea; Gastropoda) (GJ Vermeij, Ed.). PLOS ONE 11: e0156664.

