## Supplementary material for "Biofluorescence reveals hidden patterns in chitons with implications to visual ecology": File S1_Supplementary Figures

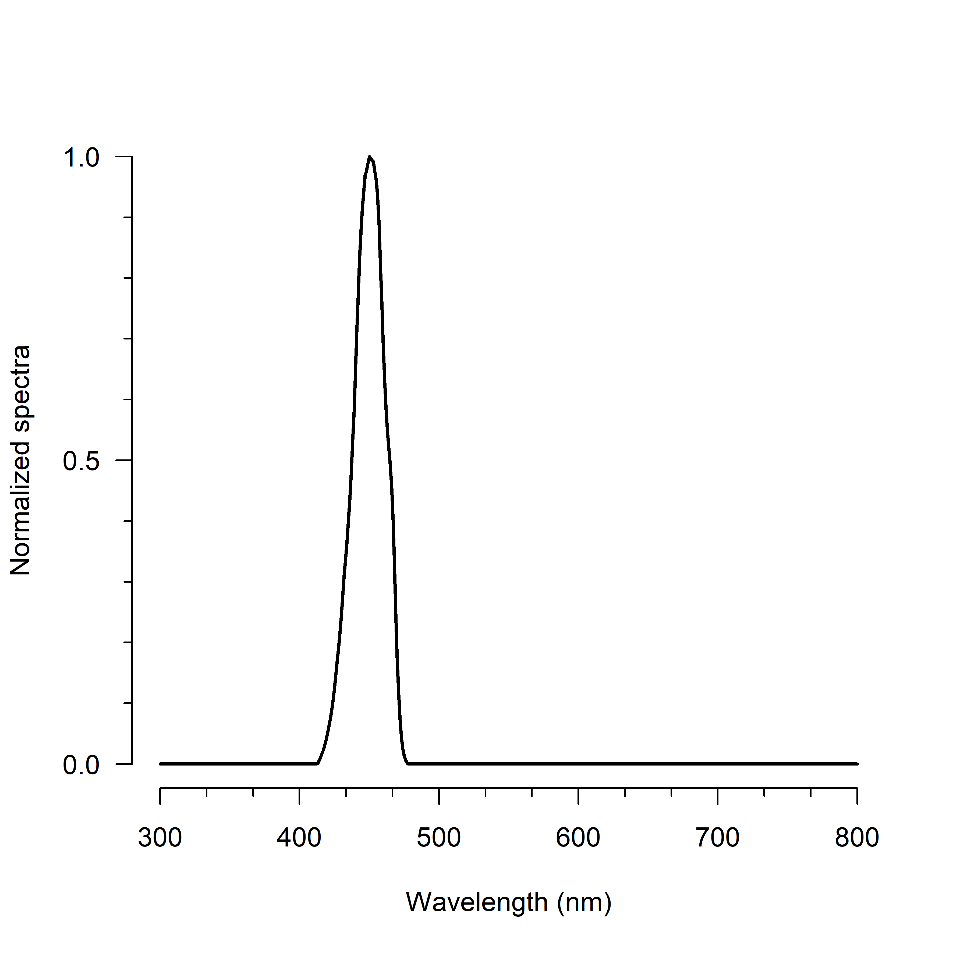

**Supplementary Figure S1:** Fluorescence excitation spectrum (µmol photon. s^-1^. m^-2^. nm^-1^).

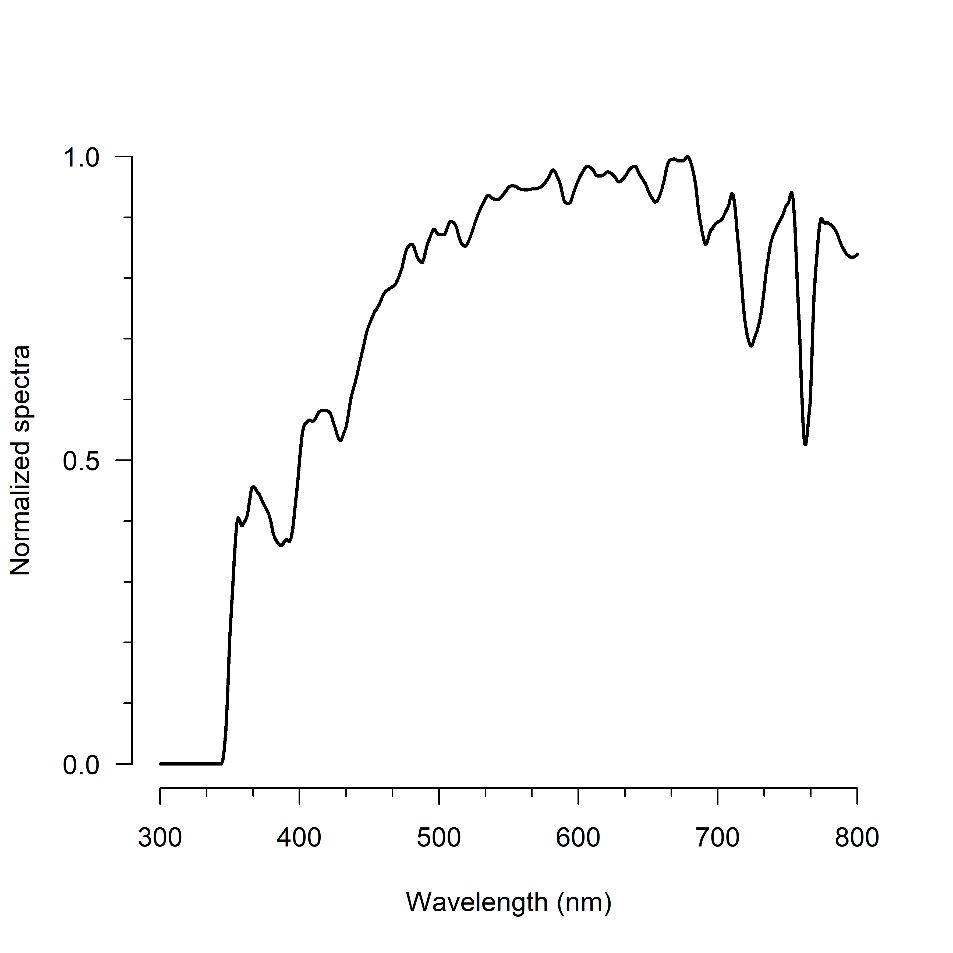

**Supplementary Figure S2**: Local irradiance spectrum (µmol photon. s^-1^. m^-2^. nm^-1^), measured outside water at 09:30am on a cloudless sunny day.

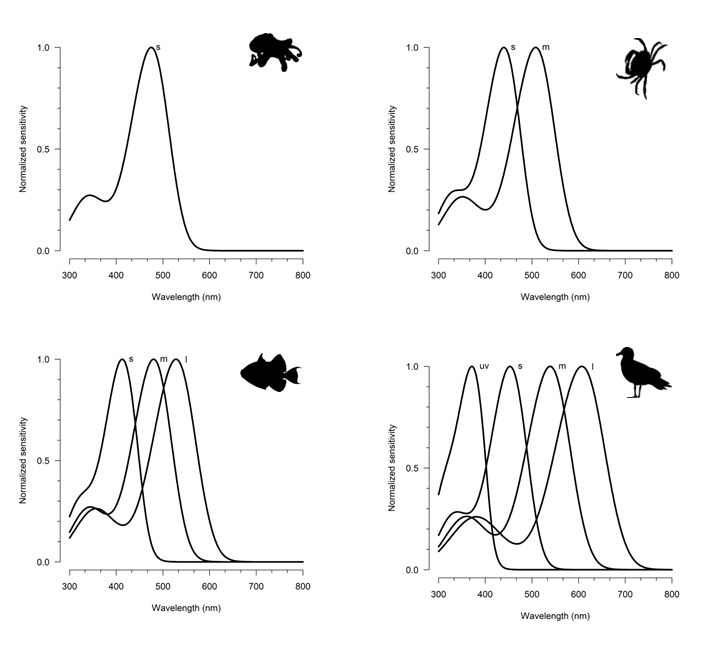

**Supplementary Figure S3:** Photoreceptor sensitivity curves from the visual systems of four potential visually guided chiton predators (i.e., octopus, shore crab, reef fish, and seagull), as documented in the literature. **Top left**: Common octopus (*Octopus vulgaris*); **Top right**: European green crab (*Carcinus maenas*); **Bottom left**: White-banded triggerfish (*Rhinecanthus aculeatus*); **Bottom right**: Blue tit (*Cyanistes caeruleus*) as a proxy to seagulls, since both have SWS1 cones of UVS type. .

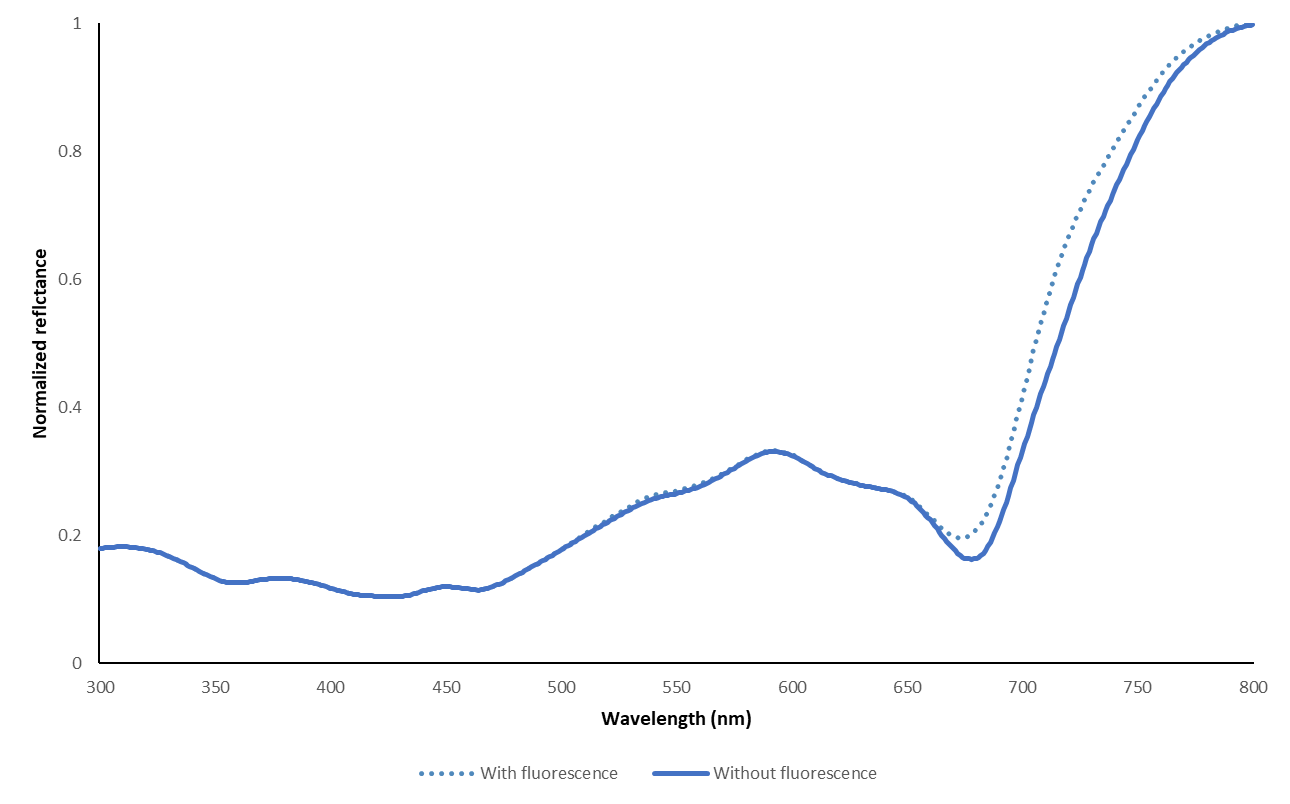

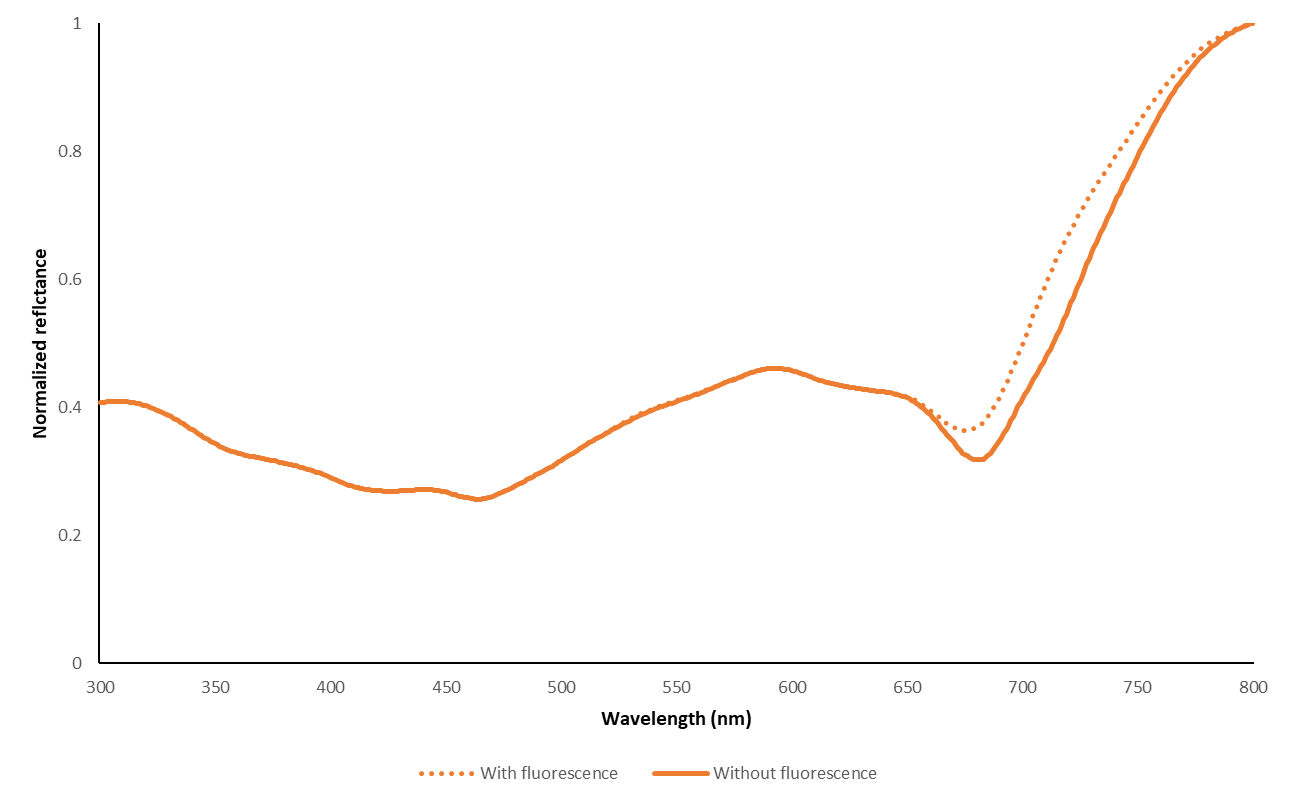

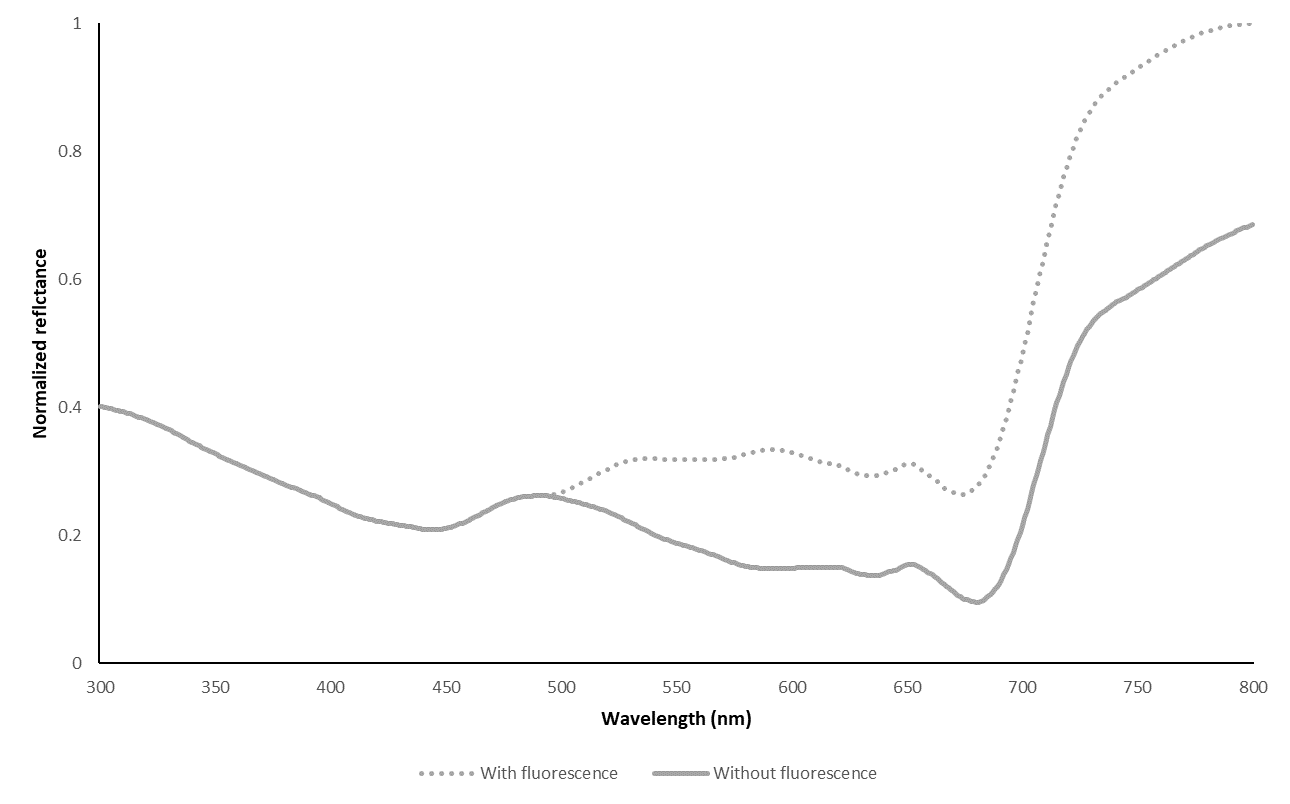

**Supplementary Figure S4**:

Comparative exitance spectra of three chitons with and without fluorescence components. *With fluorescence*: represent a combination of light reflected and fluorescence component. *Without fluorescence*: refer only to the reflected component, sometime called by the term ‘true reflectance’ (Mazel, 2017).

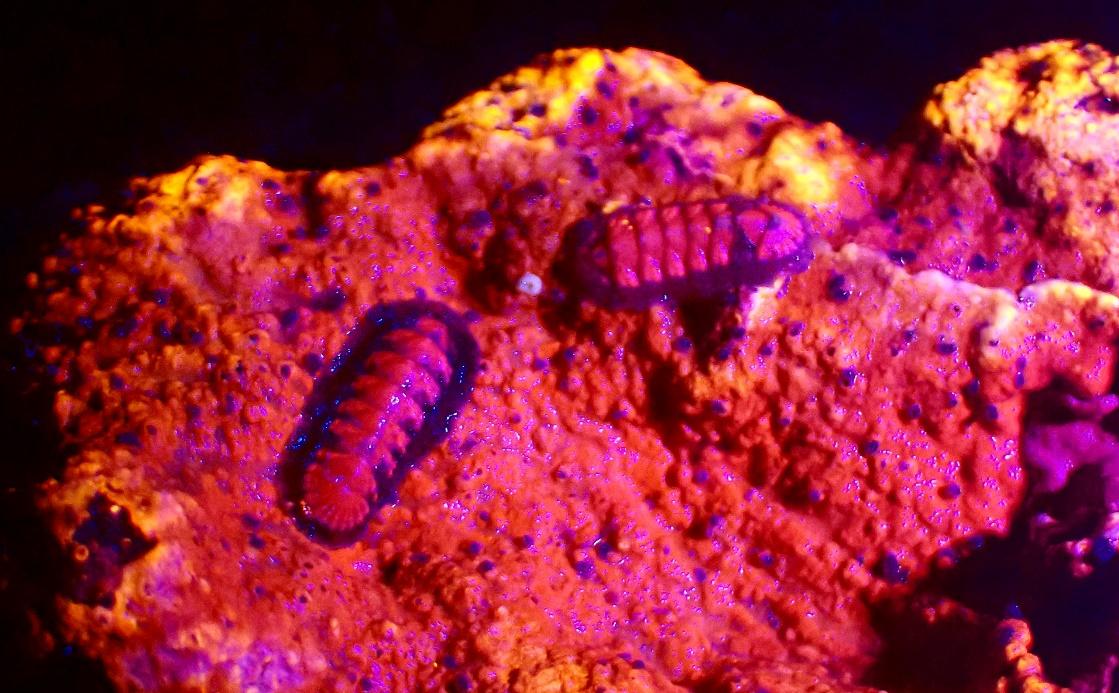

**Supplementary Figure S5**: Picture of two young individuals of *I. pectinata* attached on a rhodolith, probably in the 4th/5th stages of development, under fluorescence excitation condition.

### References

Mazel C. 2017 Method for Determining the Contribution of Fluorescence to an Optical Signature, with Implications for Postulating a Visual Function. *Front. Mar. Sci.* **4**, 266. (doi:10.3389/fmars.2017.00266)
